# Genomic characteristics and clinical significance of CD56+ Circulating Tumor Cells in Small Cell Lung Cancer

**DOI:** 10.1101/2022.04.30.487775

**Authors:** C. Ricordel, L. Chaillot, E. I. Vlachavas, M. Logotheti, A. Jouannic, T. Desvallees, G. Lecuyer, M. Aubry, G. Kontogianni, C. Mastrokalou, F. Jouan, U. Jarry, R. Corre, Y. Le Guen, T. Guillaudeux, H. Lena, A. Chatziioannou, R. Pedeux

## Abstract

**Introduction:** Circulating Tumor Cells (CTC) have been studied in various solid tumors but clinical utility of CTC in Small Cell Lung Cancer (SCLC) remains unclear. The aim of the CTC-CPC study was to develop an EpCAM-independent CTC isolation method allowing isolation of a broader range of living CTC from SCLC and decipher their genomic and biological characteristics.

**Patients and methods:** CTC-CPC is a monocentric prospective non-interventional study including treatment-naïve newly diagnosed SCLC. CD56+CTC were isolated from whole blood samples, at diagnosis and relapse after first-line treatment and submitted to Whole-exome-sequencing (WES).

**Results:** Phenotypic study confirms tumor lineage and tumorigenic properties of isolated cells for the 4 patients analyzed with WES. WES of CD56+CTC and matched tumor biopsy reveal genomic alteration frequently impaired in SCLC. At diagnosis CD56+CTC were characterized by a high mutation load, a distinct mutational profile and a unique genomic, compared to match tumors biopsies. In addition to classical pathways altered in SCLC, we found new biological processes specifically affected in CD56+CTC at diagnosis. High numeration of CD56+CTC (>7/ml) at diagnosis was associated with ES-SCLC. Comparing CD56+CTC isolated at diagnosis and relapse, we identify differentially altered oncogenic pathways (e.g. DLL3 or MAPK pathway).

**Conclusions:** We report a versatile method of CD56+CTC detection in SCLC. Numeration of CD56+CTC at diagnosis is correlated with disease extension. Isolated CD56+CTC are tumorigenic and a distinct mutational profile. We report a minimal gene set as a unique signature of CD56+CTC and identify new affected biological pathways enriched in EpCAM-independent isolated CTC in SCLC.

## Introduction

Lung cancer is the leading cause of cancer-related death worldwide, accounting for 1.6 million deaths per year (1). Small cell lung cancer (SCLC) is an aggressive subset of lung cancer, representing 10 to 15% of lung cancers and characterized by rapid-onset symptoms due to high tumor growth rate (2). At diagnosis, most patients present an extensive stage disease (ES-SCLC) for whom the recommended treatment has been for the last decades a palliative platinum-based chemotherapy regimen. Even if recently, the combination of chemotherapy with immune checkpoint inhibitors showed substantial outcomes improvement in two phase III trials (3,4), the prognosis remains poor with a median overall survival of about one year. For patients diagnosed with limited stage disease (LS-SCLC), concomitant chemo-radiotherapy is the recommended treatment option showing a median overall survival of over 20 months (5).

SCLC shows a high propensity for metastatic spreading and therefore generates a higher number of circulating tumor cells (CTC) compared to other tumor types (6). Recent studies have demonstrated that CTC isolated from patients diagnosed with SCLC can be deeply characterized at the genomic level and could have a potential prognostic significance (7–9). An in-depth analysis of single-cell CTC copy number variation (CNV) by whole-genome sequencing in a cohort of 31 SCLC patients was used to create a CNV classifier that could reliably predict chemosensitivity (10). However, these genomic analyses, especially on single cell CTC are not sustainable in routine clinical practice. Moreover, it should be noted that in most studies, CTC isolation was based on the expression of epithelial markers (EpCAM). It is well documented that tumor cells migrating through the bloodstream undergo epithelio-mesenchymal transition (EMT), characterized by the loss of expression of epithelial markers expression (11,12). Coherently, studies using a EpCAM-independent isolation method show a higher detection rate of CTC (13). A recent immunofluorescence study on a large cohort of 108 SCLC patients demonstrate that EpCAM-based methodology (Cellsearch®) was unable to detect CD56+ and/or TTF1+-CTC in more than 20% of patients (14). These results suggest that CTC isolation method relying on EpCAM markers does not circumvent the issue of EMT of CTC in SCLC. Consequently, there is a clear unmet need for comprehensive genomic studies performed on CTC isolated independently of EpCAM to achieve a better understanding of the biology of SCLC. To explore such an avenue of research, we select the CD56 marker, frequently express at surface of the membrane of SCLC cells, as it will allow the isolation of living CTC from the blood of SCLC patients.

Here, we describe a versatile and easy-to-use workflow for the detection, count and isolation of CD56+CTC from whole blood samples in a prospective cohort of 33 newly diagnosed patients with SCLC at our institution. Phenotypic analysis of isolated cells confirmed their tumor lineage. Notably, generation of CTC-Derived Xenografts (CDX) in immunocompromised mice support the tumorigenic properties of CD56+CTC. Our findings indicate that while there are genes commonly altered in biopsies and CD56+CTC samples, there is noticeable mutational diversity in the liquid biopsies, highlighting a putative distinct molecular signature. Finally, CD56+CTC count at diagnosis was associated with the stage of the disease.

## Materials and methods

### Patient and samples

We conducted a prospective non-interventional study in our institution (Rennes University Hospital), including only patients with a histologically or cytologically confirmed chemotherapy-naïve SCLC and eligible to a systemic treatment, starting from April 2016. The data lock was planned in December 2021 for all patients that achieve a minimal follow-up of at least 6 months. The recruitment is still active beyond this date. Any active neoplasia other than SCLC or known HIV, HCV or HBV infection, were considered as exclusion criterion. CD56+CTC were numerated and isolated from a whole blood sample, before the first administration of any cytotoxic agent and when disease relapsed (Fig 1A). Paired tumor tissue samples were obtained for four patients in order to compare CD56+CTC and biopsies genomic characteristics. LS-SCLC was defined as disease confined to one hemi-thorax and that could be safely encompassed in a single radiation field, otherwise patients were considered ES-SCLC. The International Association for the Study of Lung Cancer (IASCL) eighth edition of the TNM Classification for small cell lung cancer was used to stage patient’s SCLC (15). Chemosensitive status was defined as patient without progressive disease within 3 months after the first-line of treatment, as opposed to chemorefractory status. Progression-free survival (PFS) was defined as the time from the start of chemotherapy until disease progression or death from any cause and assessed using RECIST version 1.1. Overall survival (OS) was defined as the time from the start of chemotherapy until death from any cause. Patients who had not progressed at the time of the statistical analysis or who were lost to follow-up before progression or death were censored at their last evaluation. This study was conducted in accordance with the Declaration of Helsinki and was approved by Rennes University hospital ethic committee (n° 15.120). All participants provided informed consent, according French national regulation for non-interventional studies.

**Figure 1.**
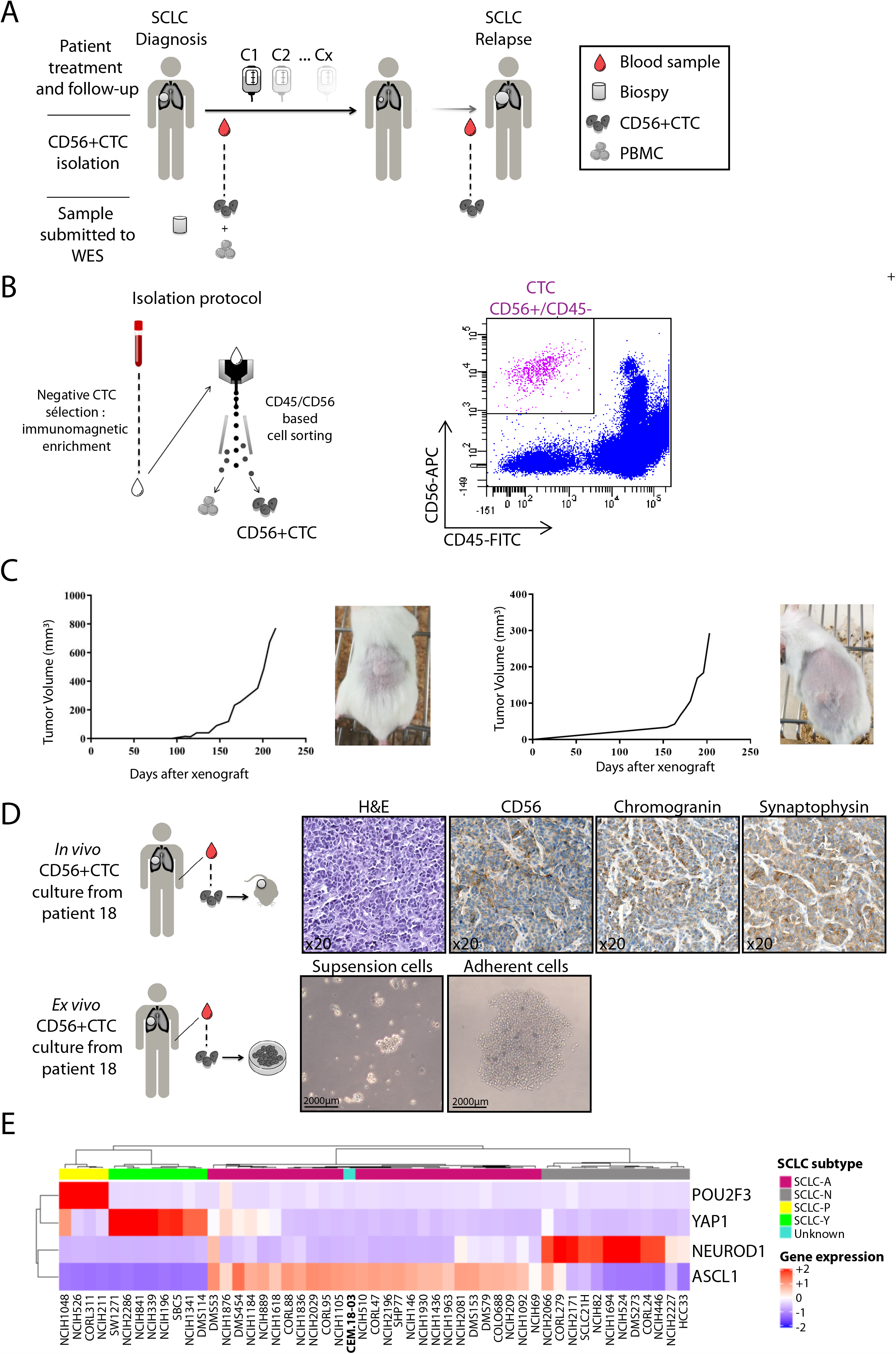
Isolation of CD56+CTCs and generation of pre-clinical models. **A**. Schematic workflow of the CTC-CPC study. **B**. Isolation procedure of CD56+CTC from whole blood sample (left panel); Representative result of flow cytometry cell sorting (right panel). **C**. Circulating tumor cells-derived xenograft (CDX) obtained from isolated CD56+CTC. **D**. IHC phenotype of *in vivo* models obtained from isolated CD56+CTC (upper panel); *Ex vivo* cell lines obtained from isolated CD56+CTC (lower panel). **E**. Heatmap and hierarchical clustering based on the expression data of the four (established) key transcription regulators that are (commonly) used to identify the SCLC molecular subtype. Samples are derived from 50 SCLC cell lines, from CCLE with known molecular subtypes and from one patient of unknown SCLC subtype (CEM 18-03). For every gene, the color scale indicates its relative expression, from blue (low) to red (high).

### Workflow Isolation of CD56+CTC from whole blood

The first step consists in a CTC enrichment from whole blood sample by immunomagnetic negative selection using the EasySep™ Direct Human CTC Enrichment Kit (StemCell Technologies®, Vancouver, Canada) according to manufacturer instruction. Because we expected to isolate a very small number of cells and to accurately sort the CTCs, the remaining cells were mixed with 1×10^5^ PBMCs (peripheral blood mononuclear cells). Cell suspension was washed in PBS 2% BSA and incubated with saturating concentrations of fluorescent[labelled antibodies against human CD56, CD45 and Cd235a (Miltenyi, Bergisch Gladbach, Germany) for 15 min at 4°C. Cells were then washed with PBS 2%-BSA and analyzed and sorted by flow cytometry using a FACSAria II (Becton Dickinson, Franklin Lakes, NJ). The population of interest was gated according to its CD56/CD45 criteria to avoid NK lymphocytes contamination (exclusion of CD56+/CD45+ cells). Cells CD56+/CD45-were sorted, centrifuged, and immediately freeze at -80°C. CD45+ PBMC were used as a negative control. Data were analyzed with FlowJo™ Software (Ashland, OR: Becton, Dickinson and Company).

### Pathway analysis and comparison of genomic mutational patterns between biopsy and paired CD56+CTC

Detailed methods for PBMC isolation, extraction of DNA, whole exome sequencing and variant annotation workflow can be found in the supplemental data.. Complete analysis was performed utilising various state-of-the-art tools. Unless stated otherwise, all bioinformatics analyses were performed using shell interface command line, on a Linux-based 64GB RAM/ 12 processor cluster server (Ubuntu 18.04). Initially, to unravel the common mutational patterns between CTCs and matched biopsies, a simple Venn diagram/overlap analysis was performed for each patient separately at the variant level (based on SNVs & INDELs) and at the gene level. The final variants that were common in the remaining variant lists of CTC and paired biopsy samples for each patient, were imported to the BioInfoMiner platform (https://bioinfominer.com) for the functional interrogation of the perturbed pathways in the CTC and biopsy samples utilizing different hierarchical and ontological vocabularies (Gene Ontology & Reactome Pathways) (16). Additionally, in order to investigate the mutational relatedness of CTC and biopsy samples, based on their mutational profiles in the six base substitution catalogues, the R package MutationalPatterns (v 1.10.0) was used (17). Finally, semantic similarity (GO functional similarity) based on the aforementioned pathway enrichment analysis was performed between CTC and biopsy samples for each patient, to identify any common molecular mechanisms.

### Derivation of an informative gene signature of CD56+CTC samples

For the derivation of a minimal gene set, expanding the molecular profile of the circulating tumor cell samples in the SCLC patients, we applied a data mining approach based on the aforementioned results of the WES computational analysis (see supplemental material and methods for details).

### In vivo xenograft experiments

NSG (*NOD*.*Cg-Prkdcscid Il2rgtm1Wjl/SzJ)* mice were purchased from Charles River Laboratories (Wilmington, MA). Mice were subcutaneously injected with a cell suspension (obtained after the first step of circulating tumor cell isolation) on the left flank with Matrigel (v/v) (Corning, Corning NY), as previously described (9). Tumor size was monitored using calliper twice per week. All animal procedures met the European Community Directive guidelines (Agreement B35-238-40 Biosit Rennes, France) and were approved by the local ethics committee (Agreement APAFIS # 7163; CEEA - 007 regional ethics committee of Brittany; France), ensuring the breed ing and the daily monitoring of the animals in the best conditions of well-being according to the law and the rule of 3R (Reduce-Refine-Replace). All experiments were performed in accordance with relevant guidelines and regulations. After tumor growth mice were euthanized (cervical dislocation) when tumors reached 1000 mm3 and tumors were harvested and cut in two pieces (one used for cell culture and the other for formalin-fixation). All experiments on live animals are reported here in accordance with ARRIVE guidelines. Refer to supplemental material & methods for RNAseq analysis and identification of SCLC patient subtype.

### Statistical analysis

Demographic and descriptive data are given as the median with the range. Categorical variables were compared with the Fisher exact test or the Pearson X^2^ test, and quantitative variables were compared with the Mann-Whitney U test when appropriate. Two-tailed P-values were reported, with P < 0.05 considered as statistically significant. The Kaplan-Meier method with log-rank test was used to perform survival analysis. Finally, binomial logistic regression model was built with the CD56^+^CTC count (□7 CD56+CTC or ≤7 CD56+CTC/ml of blood) as the dependent variable. The pre-specified cut-off of 7 CD56+CTC/ml was chosen, based on study investigating prognosis of CTC in SCLC which shows that more than 50 CTC/7.5ml (equivalent to 6.6 CTC/ml) correlate with survival (6). GraphPad Prism for Windows software (version 5.03; Graph Pad Software Inc, San Diego, CA) was used for the statistical and survival analyses.

## Results

### CD56+CTC isolated from whole blood samples are tumorigenic

To isolate CD56+CTC from SCLC patients, we have set up a two steps protocol. First, hematopoietic cells and platelets were depleted and then CD56+CTC were isolated by cell sorting, using CD56 as a positive selection marker and CD45 as a negative selection marker to avoid NK cells contamination (Fig 1B). Based on the technical validation of the isolation method (Supplementary Fig. S2A and S2B), we processed all the samples immediately after collection. A representative example of flow cytometry gating strategy used for cell sorting is shown in Figure 1B (right panel). In order to establish the tumorigenic proprieties of CD56+CTC, we performed CDX generation through implantation of isolated CD56+CTC on the flank of nude mice, according to previous published methodology (9). Out of five attempts, CD56+CTC from two patients generated tumors in mice (Figure 1C). Subsequent immunohistochemistry analysis on these CDX tumors show positivity for commonly used markers for SCLC diagnosis: CD56, synaptophysin and chromogranin A, demonstrating the stability of the patient’s neuroendocrine tumor lineage (Fig. 1D). Moreover, *in vitro* CTC-derived cell lines were obtained from two patient’s samples and amplified in culture (Fig. 1D). Consistently with the recent classification of SCLC, CDX model derived from patient 30 was successfully classified with an ASCL1 transcriptomic profile (Fig. 1E). Altogether, these results confirmed the tumorigenic properties of CD56+CTC isolated from whole blood samples of patients with a treatment-naïve SCLC.

### CD56+CTC share common genomic alterations with SCLC tissue biopsies but display a distinct mutational signature

To further characterize CD56+CTC at the genomic level, we submitted the CTC samples, matched PBMC and tumor biopsies to whole-exome-sequencing for four patients in our cohort. The average sequencing coverage (up to 135X and with a minimum depth of 20X) was 0.70, 0.87 and 0.88 for CTC, biopsy and PBMC respectively (Table 1). Many genes commonly found to be altered according to large scale genomic studies of small cell lung cancer (18), were also detected in our genomic analysis (Fig. 2A). Alterations in *TP53* or *RB1*, the two most consistently altered genes in SCLC, were found in CD56^+^CTC of 3 patients (75%). CD56+CTC from one patient didn’t show any *TP53* or *RB1* mutation but CNV analysis revealed a mono-allelic deletion of both genes (Fig. 2B). Moreover, our analysis showed that the altered pathway commonly found in SCLC, such as NOTCH signaling, were also observed in our CD56+CTC samples (Supplemental Fig. 3B). These results confirmed that the genetic profiles of CD56+CTC are similar to the known genomic landscape of SCLC.

**Figure 2.**
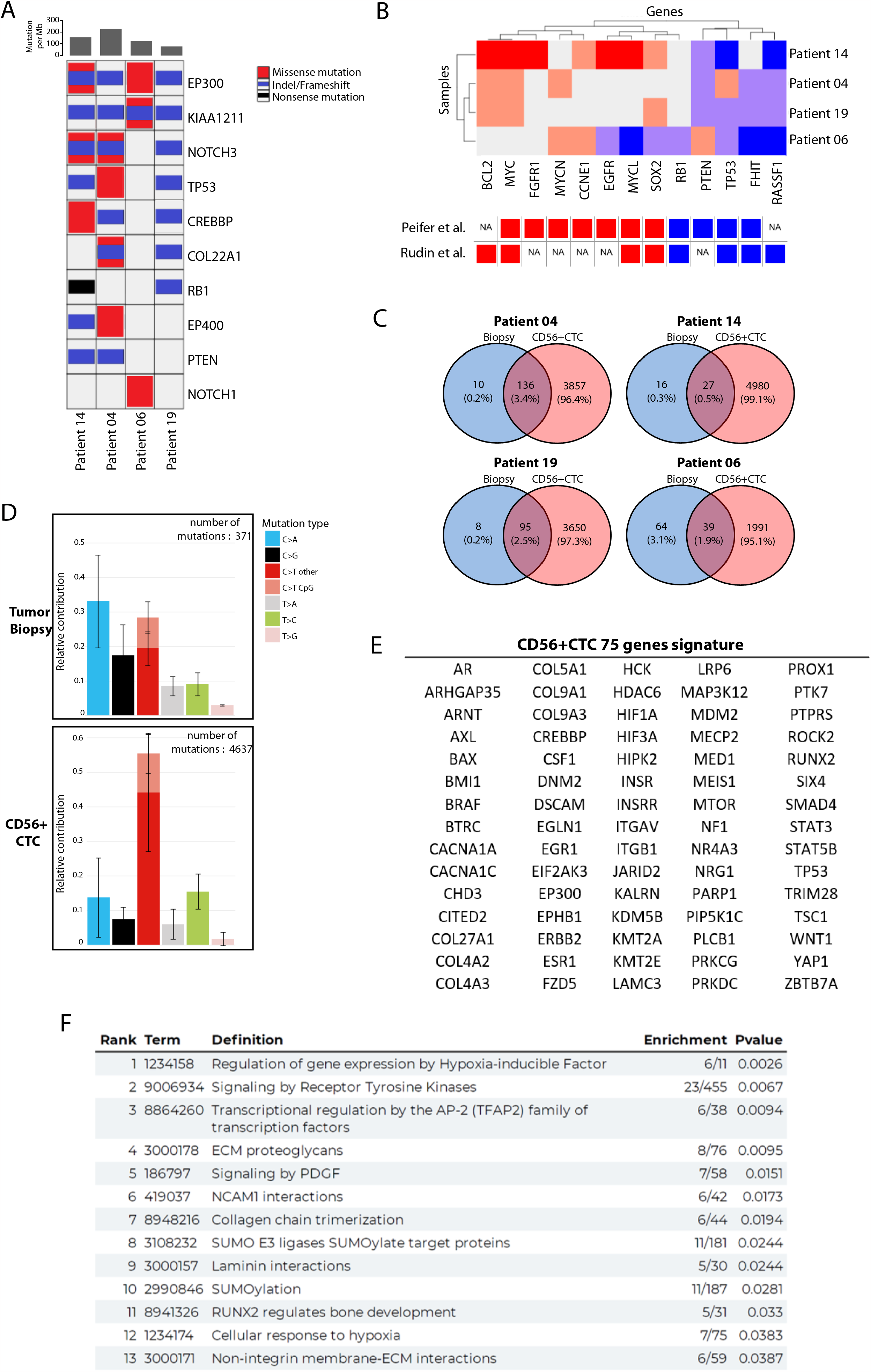
Genomic characterization of CD56+CTC. **A**. Heatmap of genomic alterations of SCLC candidate genes (CTC samples of 4 patients). Mutation rates are displayed in the top panel. **B**. Somatic copy number alterations in CD56+CTC (amplifications in red and deletions in blue); lower panel shows copy number alteration in published SCLC genomic studies. **C**. Venn diagrams represent non synonymous mutations (SNPs & InDels) in biopsies and CD56+CTCs. **D**. Mutation spectrum plot illustrates the relative contribution of the 6 base substitution types to the point mutation spectrum for each patient sample. Bars show the mean relative contribution of each mutation type over all patient samples per sample type (biopsy and CD56+CTC) and error bars indicate the standard deviation. **E**. Derived genomic signature comprised of 75 genes for the CD56+CTC at the time of diagnosis. **F**. Analysis of the Reactome pathway of the 75 genes CD56+CTC genomic signature (ranked according p-value).

Interestingly, the mutational load in CD56+CTC was much higher than in matched tumor biopsies for all patients (Table 1). The mean mutational load in CD56+CTC and tumor biopsies was 188.02 and 6.44 mutations/Mb respectively. The majority of mutations found in the tissues were confirmed in CD56+CTC (except for patient 06 were only 39/103 mutations where shared between samples) (Fig. 2C and supplemental material and method). C>A transversion, classically associated with tobacco exposure, was the dominant mutation type in the tissue biopsies (Fig. 2D). Conversely, C>T transversion (outside CpG sites) was dominant in CTC, suggesting that drivers of mutagenesis might be distinct in CD56+CTC. In line with this hypothesis, hierarchical clustering of samples based on their mutational patterns using COSMIC mutational signatures (v2-March 2015) tended to discriminate between blood and biopsy samples (Supplemental Fig.3A). Even if CTC and biopsy samples showed high similarities with multiple signatures, biopsies showed a higher correlation with signature 4 (related to tobacco mutagens), 24 & 29 (transcriptional strand bias for C>A mutations); whereas, CD56+CTC were significantly associated with signature 15 (defective DNA mismatch repair) with signatures 19 & 30 (etiology unknown), supporting the idea of distinct “circulating” drivers of mutagenesis in CD56+CTC.

To explore altered signaling pathways in CD56+CTC, we exploited BioInfoMiner (BIM) web platform which implements robust statistical and network analysis for functional enrichment investigation in order to highlight the most important biological mechanisms. Thus, we identified a genomic signature comprised of 75 hub genes for the CD56+CTC at the time of diagnosis (Fig.2E). In order to capture holistically the perturbed biological components that are related to SCLC pathophysiology and progression, we utilized one of the most comprehensive biomedical vocabularies REACTOME pathway analysis. Subsequent functional enrichment analysis of the compact gene set revealed altered biological mechanisms either directly or indirectly associated with lung cancer progression (related to neural cell adhesion, extracellular matrix organization, hypoxia or immune response) (Fig.2F). This analysis also stressed the importance of understudied processes in SCLC: sumoylation and transcriptional regulation by AP-2 and RUNX2. Subsequently, three genomic databases of SCLC (Cancer Gene Census, Intogen driver genes and MycancerGenome) were analyzed with BIM leading to the identification of 22, 17 and 9 hub genes respectively (Supplemental Table 1). Interestingly, 23 of the 75 hub genes from CD56+CTC, such as *ERBB2, TP53, CREBBP* and *NF1*, were found enriched in at least 1 of the 3 queried databases, illustrating the high affinity of the liquid biopsies to recapitulate the molecular genomic landscape of small cell lung cancer. The REACTOME pathway analysis of the 23 hub genes further confirmed that the altered biological mechanisms found in CD56+CTCs correspond to those found in SCLC, with the addition of the p53 signaling pathways (Supplemental Fig.3D).

Altogether, these observations support the hypothesis that CD56+CTC can reliably recapitulate the mutational status of treatment-naive SCLC tumors and may represent a comprehensive way of capturing the whole mutational picture of SCLC.

### Cohort demographics

From April 2016 to April 2021, 46 patients were eligible for inclusion in the study. Seven patients met the exclusion criteria and 39 patients had CD56+CTC isolation from blood samples. For five patients, isolated CD56^+^CTC were used for *in vivo*/*ex vivo* culture and one patient withdrew consent. Finally, the number of CD56+CTC at diagnosis was available for 33 patients who were therefore included in the statistical analysis (Supplementary Fig.4). The demographic and descriptive statistics of the cohort are presented in Table 2. Patients were predominantly male (69.7%), active smoker (69.7%) and had mainly an extensive-stage disease at diagnosis (66.7%). Of note, a large proportion of the patients progressed within 3 months after the end of the first-line of chemotherapy and was considered chemorefractory (45.5%). Based on the pre-specified cut-off of 7 CD56+CTC/ml, two groups of patients were compared. The two groups were well balanced with respect with age, sex, smoking exposure, chemosensitivity or median follow-up (Table 2). However, the group with more than 7 CD56+CTC/ml at diagnosis was enriched in patients with ES-SCLC (85.7% vs 52.7%), without reaching statistical significance (Fisher’s exact test; p= 0.067). Remarkably, the five patients without any metastatic localization at diagnosis show a low CD56+CTC count (mean: 3.23±3.30 cells/ml). Two patients show no detectable CD56+CTC in blood samples: one had LS-SCLC and the other one had a low thoracic tumor burden associated with multiple cerebral and medullary metastases.

### Clinical and prognosis significance of CD56+CTC

In order to correlate tumor burden and propensity to detect CTC in SCLC patients, we compared the number of CD56+CTC according to the initial stage of the disease. Patients with ES-SCLC show significantly higher number of CD56+CTC compared with patient diagnosed with LS-SCLC (respective median of 7.95 vs 2.00; p=0.014) (Fig. 3A). To gain further insight into the specific clinical parameters associated with higher numbers of CTCs detected, we looked at each descriptor of the TNM classification (Fig 3B). Even if we observed a trend toward a higher count of CD56+CTC in patient with more advanced T, N or M status, statistical significance was not reached. As of December 2021, a relapse or death event occurred in all patients in our cohort after first line-treatment. The median PFS was 5.2 months in the overall cohort with 5.2 months and 5.3 months in the low CD56+CTC group and high CD56+CTC group, respectively (HR=0.80 CI 95% [0.39-1.56]; p=0.5) (figure 3C, left panel). Median overall survival was 8.1 months in the overall cohort and no statistically significant difference was observed between the two groups: 10.16 months in the low CD56+CTC group versus 8.7 months in the high CD56^+^CTC group (HR=0.92 CI 95% [0.43-1.93]; p=0.8) (figure 3C, right panel). Importantly, at the time of analysis, no difference in survival was observed between ES-SCLC and LS-SCLC in our cohort (Supplemental Fig.5).

**Figure 3.**
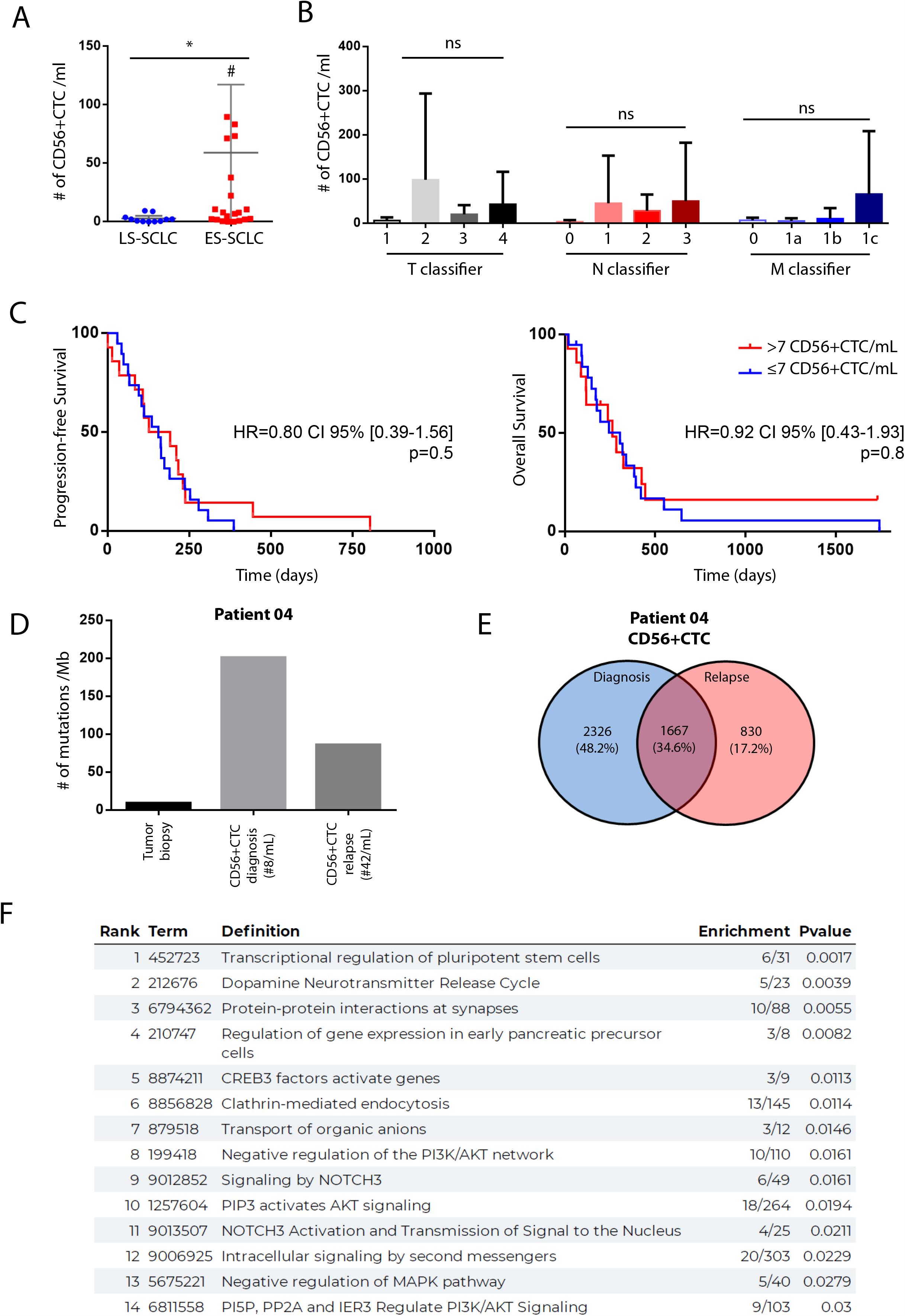
Clinical significance of CD56+CTC at diagnosis and somatic evolution of CTC at relapse. **A**. Diagnosis numeration of CD56+CTC according to SCLC staging (*: p<0.05; # : two values outside the limits of the graph). **B**. Numeration of CD56+CTC according to the 8^th^ TNM IASCL classification (ns : non-significant) **C**. Kaplan-Meier curve of progression-free-survival (left panel) or overall survival (right panel) according to the numeration of CD56+CTC with a pre-specified cut-off value of 7 CTC/ml. **D**. Mutational load of all samples from patient 04. **R**. Venn diagrams representing non synonymous mutations (SNPs & InDels) in CTCs at diagnosis and after relapse (right) for patient 04. **F**. Analysis of the REACTOME pathways of the common non synonymous genes (1667) at diagnosis and after relapse.

### Somatic evolution of CD56+CTC at progression after first-line treatment

In order to decipher the tumor biology of SCLC at relapse, we conducted CD56+CTC isolation for patient 04 at clinical progression (costal metastasis after 3 cycles of carboplatin-etoposide). CD56+CTC counts were higher at relapse compared to diagnosis: 42 cells/ml versus 8 cells/ml respectively. However, WES showed that the mutation load decreased almost two fold in CD56+CTC at relapse (Fig. 3D). Interestingly, most of the mutations (1667/2497=66.7%) observed in relapsed CD56+CTC were also detected in CD56+CTC isolated at diagnosis (Fig. 3E). Conducting functional analysis enrichment, we were able to identify differentially altered pathways between diagnosis and relapse circulating CD56+ tumor clones (Fig. 3F). Notably, signaling pathways known to drive tumorigenesis in SCLC and currently investigated in targeted therapy trials (DLL3-targeting molecules for the NOTCH pathway or PI3KCA/AKT inhibitors for the MAPK pathway) were specifically implicated in relapsed CD56+CTC (19,20). These results suggest that overcoming relapse after first-line systemic treatment may require targeting these signaling pathways, as we speculate this gives a selective advantage for resurgent SCLC clones.

## Discussion

This prospective study demonstrates the feasibility of an EpCAM-independent and immunofluorescence-based CTC isolation in SCLC and its application in clinical practice. We confirmed that CD56+CTC display a neuroendocrine tumor lineage and conserve tumorigenic properties *in vivo* and *in vitro*. Coherently, whole-exome sequencing validated that CD56+CTC show typical genomic alterations and classical altered pathways commonly found in SCLC (e.g. TP53 mutation, RB1 loss) but they are also characterized by distinct mutational patterns. We prospectively observed that ES-SCLC display a higher CD56+CTC count at diagnosis compared to LS-SCLC. Finally, we show that detection of CD56+CTC at progression after first-line treatment uncover new oncogenic pathways implicated in relapse circulating tumor cells.

In line with previous results in non-small cell lung cancer, EpCAM-independent detection of CTC show a high sensitivity (21). Indeed, median number of CTCs at baseline using CellSearch® approach in a large SCLC cohort was 14 CTC/7.5ml of blood (14), compared to 28 CTC/7.5ml of blood in our cohort. Interestingly, in a recent cohort restricted to ES-SCLC, the median number of CTC at diagnosis using CellSearch® was 30 CTC/7.5 ml of blood (22), whereas it reaches 225 CTC/7.5 ml of blood in the ES-SCLC patients in our cohort. As EMT is expected to be a more potent phenomenon when metastatic spread occurs, we speculate that our EpCAM-independent method allow us to capture CTCs from SCLC that would remain undetected using CellSearch® approach, especially in late-stage SCLC. Supporting this hypothesis, similar observations were made in a large cohort of 108 SCLC patients where CellSearch® failed to detect CTCs for 22 patients whereas immunofluorescence-based approach succeeded (14). Moreover, intra- and inter-patient heterogeneity was observed for EMT markers in both CTCs and circulating tumor microemboli in lung cancer (23) and could partially explain these discrepancies.

To our knowledge, this study is the first to report genomic features of EpCAM-independent CTC in SCLC. Beyond the confirmation of typical genomic alteration of SCLC in CD56+CTC, we show that the mutational load was high compared to paired tumor biopsies or published data on CTC of SCLC. This suggests that our EpCAM-independent method allows us to capture more exhaustively CTC heterogeneity and altered signaling pathways. To support this hypothesis we report a set of 75 hub genes altered in CD56+CTC, comprising 23 core genes shared with public SCLC genomic databases. Unique CTC genomic signatures have been described in others solid malignancies (24,25), highlighting cell populations with different functional or metastatic potential. In the context of SCLC, larger genomic studies considering single-cell sequencing of EpCAM-independent CTC are needed to confirm the new pathways we identified as altered in our dataset. Further longitudinal data from sequencing of CTC at diagnosis and relapse are underway to establish firmly the relevant common mechanisms that might drive resistance to therapy.

Prognostic value of CTC from SCLC has been extensively reported before. Indeed, Hou et al. reported that more than 50 CTCs/7.5 ml of blood at diagnosis has a detrimental effect on PFS and OS (6). Similarly, Su et al. and Carter et al. reported that copy-number-alteration (CNA) analysis from single cell CTC could predict chemosensitity status and PFS (7,10). In our study, no difference of CD56+CTC numeration was observed based on the chemosensitivity status of tumors and no association with prognosis was found based on the pre-specified cut-off value of 7 CD56+CTC/ml. In line with this result, a recent work from Masseratakis et al. et using an immunofluorescence CTC-isolation method did not show correlation between CD56+CTC and clinical outcomes (14). Altogether, we hypothesized that CD56+CTC might not correlate with the invasive potential or the aggressiveness of the primitive tumor and/or might not dictate tumor sensitivity to systemic therapy. One caveat to this hypothesis is the absence of an impact on survival of the stage of the disease in our cohort. Some confounding factors and the relative small size of our cohort might have limited the power of the survival analysis. A better understanding of the biology of EpCAM-negative CTC and of the processes of EMT in SCLC is needed to definitely address this question.

Our study has several limitations. The cohort is relatively small and patients were recruited in a single institution. A direct comparison of EpCAM-independent isolation method and CellSearch® could not be performed as this technology is not available in our institution. We also acknowledge that using CD56 as positive CTC selection marker might preferentially capture CTC retaining neuroendocrine phenotype. However, it is unclear today if EMT impacts neuroendocrine state of CTC. Masseratakis et al, confirmed the favorable comparison of an immunofluorescence-based CTC isolation from SCLC (using CD56 and/or TTF1) compared to the CellSearch® technology (14). However, no study has explored to date the potential co-expression between EpCAM and CD56 markers on CTC from SCLC. Of note, as combination of anti PD-L1 with chemotherapy was not at a standard of care during the larger inclusion period of the study, we couldn’t assess the relevance of CD56+CTC upon immunotherapy. However, as EMT has been associated with inflammatory tumor microenvironment and elevation of multiple targetable immune checkpoint molecules in lung cancer (26), exploring the relation between CD56+CTC and EMT in the context of the new immunotherapeutic strategies in SCLC would be an important avenue to explore.

## Conclusion

Isolation of CD56+CTC using an immunofluorescence-based EpCAM-independent method at diagnosis is feasible in SCLC and might capture more efficiently tumor genomic heterogeneity. CD56+CTC are characterized by a high mutation load and a distinct mutational signature from paired tumor biopsies. Above classical pathways altered in SCLC, we identify new biological processes specifically affected in CD56+CTC. High numeration of CD56+CTC is associated with ES-SCLC. Finally, detection of CD56+CTC at progression after first-line treatment might help to understand somatic evolution of SCLC and elaborate strategies overcoming therapeutic resistance.

## Figure and tables legends

**Table 1. Genomic analysis of tumor biopsies and CD56+CTC.**

**Table 2. Cohort demographics.**

## Acknowledgement

We’d like to thank, la Region Bretagne, La Ligue Contre le Cancer (Grand Ouest), l’Association pour la Recherche sur le Cancer (ARC) and Roche® company for their financial supports to the CTC-CPC study. This work was also supported by ANR through the Labcom Oncotrial projet 2014–2019 (Université de Rennes 1 – UAR Biosit – BIOTRIAL, Rennes, France), notably through grants to UJ, RP & TG, and by the “Région Bretagne” as part of a collaborative project with Biotrial supported by “Biotech Santé Bretagne”. We thank the cell sorting facility (Cytometry, UAR Biosit, Rennes, France), the histopathology facility (H2P2, UAR Biosit, Rennes, France) for their technical expertise and the animal facility the animal facility (Arche, UAR Biosit, Rennes, France) for animal husbandry and care. We thank the DRCI department of the CHU Rennes for administrative support.

## Data and code availability

Whole-exome sequencing data have been deposited in the ArrayExpress database at EMBL-EBI (www.ebi.ac.uk/arrayexpress) under accession number E-MTAB-10766 at eh following address : https://www.ebi.ac.uk/biostudies/arrayexpress/studies/E-MTAB-10766?key=362b5948-a5e9-4026-bbef-943c1703cbdf. Additional supplementary files from the computational integrative workflow, R scripts and command line code implemented to reproduce the analysis are available on the Zenodo open data repository

## Notes

**Conflict of interest statement:** CR declare personal and institutional financial interest as invited speaker for Takeda, Astrazeneca, as advisory role for BMS, MSD and Takeda; LC declare no conflict of interest; EIV declare no conflict of interest; ML declare no conflict of interest; AJ declare no conflict of interest; TD declare no conflict of interest; GL declare no conflict of interest; MA declare no conflict of interest; GK declare no conflict of interest; CM declare no conflict of interest; FJ declare no conflict of interest; UJ declare no conflict of interest; RC personal interest as invited speaker for BMS, as consulting or advisory role for Takeda, Lilly, Sanofi and Astrazeneca; YLG declare no conflict of interest; TG declare personal and institutional financial interest as member of the board of directors of Kineta; HL declare personal financial interest as honoraria recipient for BMS, Astrazeneca, MSD, Pfizer, Sandoz, as consulting or advisory role for Astrazeneca, Pfizer, Sanofi, Daiichi Sankyo/Lilly, Novartis and Roche; AC declare personal and institutional financial interest as CEO of e-NIOS Applications PC; RP declare no conflict of interest; CTC-CPC study was initially funded by the JOTGO 2015 research grant with CR as a recipient.

**Sources of support :** Région Bretagne, La Ligue Contre le Cancer (Grand Ouest), Association pour la Recherche sur le Cancer (ARC), ANR through the Labcom Oncotrial projet 2014–2019 (Université de Rennes 1 – UAR Biosit – BIOTRIAL, Rennes, France) and JOTGO 2015 grant from Roche® company.

### Competing Interest Statement

The authors have declared no competing interest.

### Summary of Updates

Minor edits on Figure 1 Details adedd in material and methods section

