## Supplemental information for "Genomic characteristics and clinical significance of CD56+ Circulating Tumor Cells in Small Cell Lung Cancer"

- Supplemental Figure and Table legends:

**Supplemental Figure 1. Schematic illustration of the Variant Annotation workflow employed in this study.** The multi-step analysis procedure includes elimination of variants with low quality reads, variant annotation, filtering of the genetic variants based on the effect prediction, as well as functional and frequency filtering of the final lists of variants. The final lists of genes from each sample occurred from the indels and rare SNPs heating exonic and splice site regions. The workflow also provides information concerning the somatic status in lung cancer datasets that are recorded in COSMIC database and the occurrence, which indicates the relative number of the recorded COSMIC samples for the resulted variants.

**Supplemental Figure 2. Efficiency of the EpCAM-independent immunofluorescence-based CD56**+**CTC isolation method.** **A.** Representative images of flow cytometry plot (upper panel) with indicated number of NCI-H69 cells contaminating 20ml of whole blood sample. Results are represented in a column graph with standard deviation (lower panel; n=3). **B.** Representative images of flow cytometry plot (upper panel) after indicated lapse time of contamination of 20ml of whole blood sample with 5000 NCI-H69 cells. Results are represented in a column graph with standard deviation (lower panel; n=3).

**Supplemental Figure 3. Pathways altered in CD56**+**CTC and mutational signature. A**. Heatmap of pairwise cosine similarity between the mutational profile of each sample and available COSMIC signatures. The signatures have been ordered according to hierarchical clustering (average linkage) using the cosine similarity between signatures, such that similar signatures are displayed close together. **B.** Signaling pathways recurrently affected and found significantly enriched after pathway analysis with cBioPortal in CD56+CTC. Red and blue boxes denote genes with activating and inactivating alterations, respectively. **C**. Derived genomic signature comprised of 75 genes for the CD56+CTC at the time of diagnosis (left panel). The respective color denotes the relative overlap of the 75 genes with 3 clinical databases (Cancer Gene Census, Intogen driver genes and MycancerGenome; red:≥1/3 incidence, blue:3/3 incidence) and are summarized in the derived genomic signature comprised of 23 hub genes for the CD56+CTC at the time of diagnosis (right panel)

**D.** Analysis of the Reactome pathway of the 23 genes hub CTCs genomic signature.

**Supplemental Figure 4. Flow chart** of CTC-CPC study (data cut-off : 31/12/21)

**Supplemental Figure 5. Kaplan-Meier curve of overall survival** according to the stage of the SCLC (ES: extensive-stage; LS : limited-stage).

**Supplemental Table 1.** CD56+CTC gene signature compared to SCLC tissue database. The respective color denotes the relative overlap of the 75 genes with 3 clinical databases (Cancer Gene Census, Intogen driver genes and MycancerGenome)

- Supplemental material and method :

*PBMC isolation from whole blood samples*

The blood sample collected in EDTA-K2 tube (1x5mL tubes) was diluted in an equal volume of PBS 2% BSA. Five milliters of Ficoll-Paque™ (Miltenyi) was carefully added on the suspension, and then centrifuged 20 minutes at 1200 x g at room temperature without brake. The mononuclear cell layer was carefully aspirated and diluted in an equal volume of PBS 2% BSA. The suspension was centrifuged, washed in PBS 2% BSA and immediately freeze at -80°C.

*H-69 cells count from whole blood samples*

For experimental development of our CTC isolation procedure, blood samples from healthy donors were collected in heparin tubes (2x10mL tubes) and mixed with 5000 to 750 H-69 cells and then processed at different time point (H0, H24, H48) (see Supplemental Figure S1A and S1B). Approximately 10 ml of the heparin blood samples were gently mixed and incubated with a CTC enrichiment coktail (StemCell Technologies®) at room temperature for 5 min in constant mixing. After dilution of 1:2 in PBS 2%-BSA the blood was incubated with buffer EasySep™ and RapidSpheres™ in “Big Easy” StemCell magnet for 10 minutes. The cells suspension was collected and the same immunomagnetic depletion step was repeated once. Then cells suspension was centrifuged and the pellet resuspended in 70µL of PBS 2% BSA. Because we expected to isolate a very small number of cells and to accurately sort the CTCs, the remaining cells were mixed with 1x10^5^ PBMCs (peripheral blood mononuclear cells). Cell suspension was washed in PBS 2% BSA and incubated with saturating concentrations of fluorescent‐labelled antibodies against human CD56, CD45 and Cd235a (Miltenyi, Bergisch Gladbach, Germany) for 15 min at 4°C. Cells were then washed with PBS 2%-BSA and analyzed by flow cytometry using a NovoCyte cytometer (Agilent technologies, CA). The population of interest was gated according to its CD56/CD45 criteria to avoid NK lymphocytes contamination (exclusion of CD56+/CD45+ cells). Cells CD56+/CD45- were counted and CD45+ PBMC were used as a negative control. Data were analyzed with NovoExpress 1.2.5 Software (ACEA Biosciences, CA).

*CTC enrichment from whole blood samples*

Blood samples were collected in EDTA-K2 tube (1x5mL tube) and in heparin tube (2x10mL tubes) before first chemotherapy administration. Approximately 10 ml of the heparin blood samples were gently mixed and incubated with a CTC enrichiment coktail (StemCell Technologies®) at room temperature for 5 min in constant mixing. After dilution of 1:2 in PBS 2%-BSA the blood was incubated with buffer EasySep™ and RapidSpheres™ in “Big Easy” StemCell magnet for 10 minutes. The cells suspension was collected and the same immunomagnetic depletion step was repeated once. Then cells suspension was centrifuged and the pellet resuspended in 70µL of PBS 2% BSA.

*Cell culture*

Tumors were dissociated using the human tumor dissociation kit (Miltenyi) associated to the gentle MACS dissociator (Miltenyi Biotec, Paris) according to manufacturer's recommendations and cells were directly cultured in DMEM medium (Gibco, EnglandThermoFisher Scientific, Waltham, MA) supplemented with 5% fetal bovine serum (SVF), 0.5% Penicillin-Streptomycin, 8,4 ng/mL cholera toxin, 5ug/mL insulin, 0,4 ug/mL hydrocortisone, 24 ug/mL adenine and 10 ng/mL EGF.

*Immunohistochemistry*

Five-micron sections of formalin-fixed paraffin-embedded tissues were transferred to glass slides and incubated in TBST in the presence of 5% BSA. Immunostaining was performed using an automated slide staining system (Discovery XT -Ventana Medical Systems). Whole slide image acquisition was performed using a Nanozoomer 2.0-HT software (Hamamatsu Photonics K.K., Hamamatsu, Japan). Antibodies against human CD56 (123C3.D5), Synaptophysine and Chromogramine A Ab-3 (clone LK2H10 PHE5) were obtained from ThermoFischer Scientific.

*Extraction of DNA*

Genomic DNA (DNAg) of CTC and PBMC were prepared using the REPLI-g Single Cell Kit and QIAamp DNA Mini Kit respectively (both from Quiagen, Hilden, Germany), according to manufacturer's recommendations. Tumor biopsies were obtained from primary site for 4 patients. Biopsy sections was deparaffinized with Deparaffinization Solution (Qiagen) at 56°C for 3 minutes. Biopsy DNA was extracted using QIAamp DNA FFPE Tissue Kit (Quiagen) according to manufacturer's recommendations.

*Whole exome sequencing*

For each sample, total genomic DNA library was prepared and control quality of process (Fragment Analyzer and qPCR assays) was performed at each step. Library preparation, exome capture and sequencing were performed by Integragen SA (Evry, France) according to manufacturer’s instruction and relative protocols (SureSelect, Agilent) without modification except for library preparation performed with NEBNext® Ultra II kit (New England Biolabs®). Genomic DNA was captured using Agilent in-solution enrichment methodology with their biotinylated oligonucleotides probes library (SureSelect Clinical Research Exome V2, Agilent Technologies). Sequencing was performed on a Illumina® HiSeq4000™ platform in paired-end mode (2x75 bases). The average depth of constitutional, CTC and tumor exomes were 80X, 135X and 135X, respectively. Whole-exome sequencing data have been deposited in the ArrayExpress database at EMBL-EBI (www.ebi.ac.uk/arrayexpress) under accession number E-MTAB-10766.

Somatic variant discovery

The raw sequencing reads were preprocessed through initial quality control with FastQC (https://www.bioinformatics.babraham.ac.uk/projects/fastqc/). Next, the resulting reads were aligned to the hg38 reference genome with the Burrows-Wheeler software package (BWA-MEM 0.7.12-r1039) (Li & Durbin, 2009). Somatic variant discovery (both somatic point mutations and insertions/deletions) followed through the updated GATK4 Mutect2 (4.1.3) pipeline (McKenna et al., 2010). Briefly, the workflow included base quality score recalibration, indel re-alignment, duplicate masking and SNP/INDEL discovery across tumor-normal paired samples simultaneously using additional filtering parameters, as suggested by the GATK4 Best Practices recommendations. In parallel, somatic copy number alteration analysis was performed using the respective GATK4 CNV workflow (https://gatk.broadinstitute.org/hc/en-us/articles/360035531092?id=11682). Briefly, the pipeline includes a multitude of levers, based on denoising case sample alignment data against a created panel of normals (PoN-matched normal samples in our analysis) to obtain copy ratios, modeling and grouping continuous copy ratios into segments based on a Gaussian-Kernel binary segmentation algorithm, and finally the characterization of neutral, amplification and deletion events for the segmented copy ratios. The resulted copy ratio alterations were used as an input to the R package CNVRanger (v1.0.3), to perform downstream analysis. Firstly, the summarization of the individual CNV calls across the CTC patients into summarized regions, is comprised of two main steps: trimming of low-density areas (regional density less than 10% of the number of calls within a summarized region) with CNVRuler, and identification of recurrent CNV regions (frequency more than one of the 4 CTC samples) based on the GISTIC algorithm. Finally, the findOverlaps function from the R package GenomicRanges was used to identify the protein-coding genes overlapping the summarized CNV regions.

Variant Annotation workflow

After variant calling, the somatic mutations were annotated to the corresponding genes and then filtered, based on a workflow created and implemented in Seven Bridges Genomics (https://www.sevenbridges.com) as shown in Supplemental Figure 1. In the first step of the workflow the variants were subjected to an initial pre-filtering process. More specifically, variants with a filter flag other than PASS were excluded and variants with biallelic sites were kept. The second step included genomic annotation, as well as filtering of genetic variants based on the effect prediction. The selected effects of Single Nucleotide Polymorphisms (SNPs) and short Insertions and Deletions (InDels) in exonic regions included missense variants, non-sense variants, stop lost variants, start gained variants, start lost variants, frameshift InDels, and in splice site regions: splice region variants, splice acceptor variants and splice donor variants. The last step included functional and frequency filtering. The frequency filtering included exclusion of variants with minor allele frequency > 0.01 in widely used databases. Based on dbSNP build150 database (http://varianttools.sourceforge.net/Annotation/DbSNP), SNPs that have already been recorded with the corresponding rs identifier were reported. Functional filtering was also performed for SNPs. More specifically, variants based on scores resulting from CADD (Combined Annotation Dependent Depletion) database were included in the analysis. CADD score integrates the information from many various functional annotations and condenses this information into a single score. The minimum CADD threshold in the specific case was set to 20. Information concerning the somatic status in lung cancer datasets that are recorded in COSMIC database (https://cancer.sanger.ac.uk/cosmic) and the occurrence which indicates the number of the recorded COSMIC samples for the resulted variants were included in the resulted output. The heatmaps plots of genomic data were generated with oncoPrint function from ComplexHeatmap R package (December 2022) :

https://github.com/jokergoo/ComplexHeatmap/blob/master/R/oncoPrint.

*Derivation of an informative gene signature of CD56+CTC samples*

For the derivation of a minimal gene set, expanding the molecular profile of the circulating tumor cell samples in the SCLC patients, we applied a data mining approach based on the aforementioned results of the WES computational analysis. Briefly, the first step included the integration of two distinct gene lists. The one, included a subset of recurrent somatic alterations in the CTC samples, which were present in at least in three out of the four patients (at the gene level, union of SNVs & InDels), whereas the other included the union of the copy number alterations (gain or loss), as described in Somatic variants discovery section (see supplementals). The final merged gene list, was then further filtered using an independent microarray dataset of human SCLC cell lines ^2^. The raw microarray data were analyzed through R/Bioconductor software (R version 3.6.1) ^3,4^. Normalization and pre-processing were performed using the R package *affycoretools* (v 1.56.0) ^5^. Additionally, to remove unexpressed genes, we applied a non-specific intensity filtering based on the density of the probeset expression values, to remove genes that were unexpressed in less than 20% of the total samples. It resulted in 2343 unique genes, representing the total genomic alterations of the CTC samples. Next, an additional dimensionality reduction of the merged feature set was conducted by selecting the “altered genes”, which were also identified as “expressed” based on various SCLC cell lines gene expression microarray dataset (9731 genes). The resulting gene list, was used as an input to the BioInfoMiner web platform, resulted in a final set of 75 hub genes. A subsequent query of this signature to various cancer genomics databases and data portals-oriented to SCLC highlighted specific gene components, with pivotal role to lung cancer progression and manifestation. In parallel, we compared the final signature with the cBioPortal for Cancer Genomics open-source resource, utilizing 3 available SCLC studies (Cancer Gene Census, Intogen driver genes and MycancerGenome) with 229 patients in total.

RNAseq analysis and identification of SCLC patient subtype

Recent studies showed that SCLC subtypes can be defined by the differential expression of four key transcription regulators: achaete-scute homologue 1 (ASCL1; also known as ASH1), neurogenic differentiation factor 1 (NeuroD1), yes-associated protein 1 (YAP1) and POU class 2 homeobox 3 (POU2F3). The molecular subtype of one selected patient with available gene expression data was therefore determined by performing hierarchical clustering of that patient with the 50 cell lines of known SCLC type available from the Cancer Cell Line Encyclopedia (CCLE). Briefly, RNA-seq raw data from the patient was processed with Cutadapt and FASTQC for the trimming and quality control, respectively. Reads were aligned to the hg38 reference genome with STAR and the genes relative abundance levels were quantified with RSEM software package using the v33 gene annotation file from GENCODE. Patient data and RNA-seq gene expression data from the 50 cell lines of known molecular subtypes (also obtained with RSEM) were pooled. We calculated TPM (Transcripts Per Million) and performed TMM (Trimmed Mean of M-values) normalization and scaling using the EdgeR R package. The heatmap plot was created with the ComplexHeatmap R package using the euclidean distance and the Ward.D2 linkage method.

- Supplemental information references:
