## Supplementary figures and images for "Genomic characteristics and clinical significance of CD56+ Circulating Tumor Cells in Small Cell Lung Cancer"

### Supplemental Fig.1

Supplemental Fig1

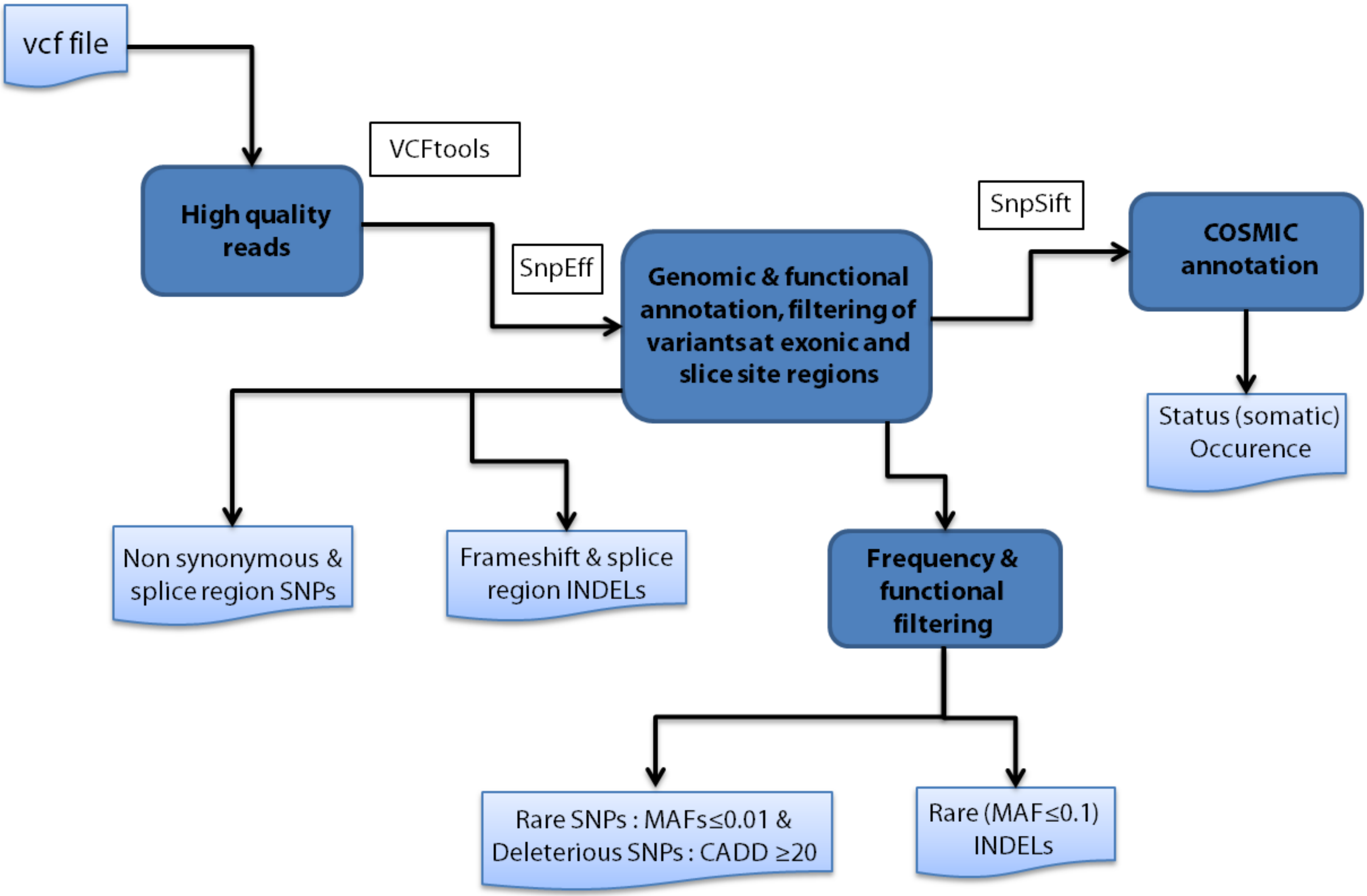

### Supplemental Fig.2

#### Supplemental Fig2

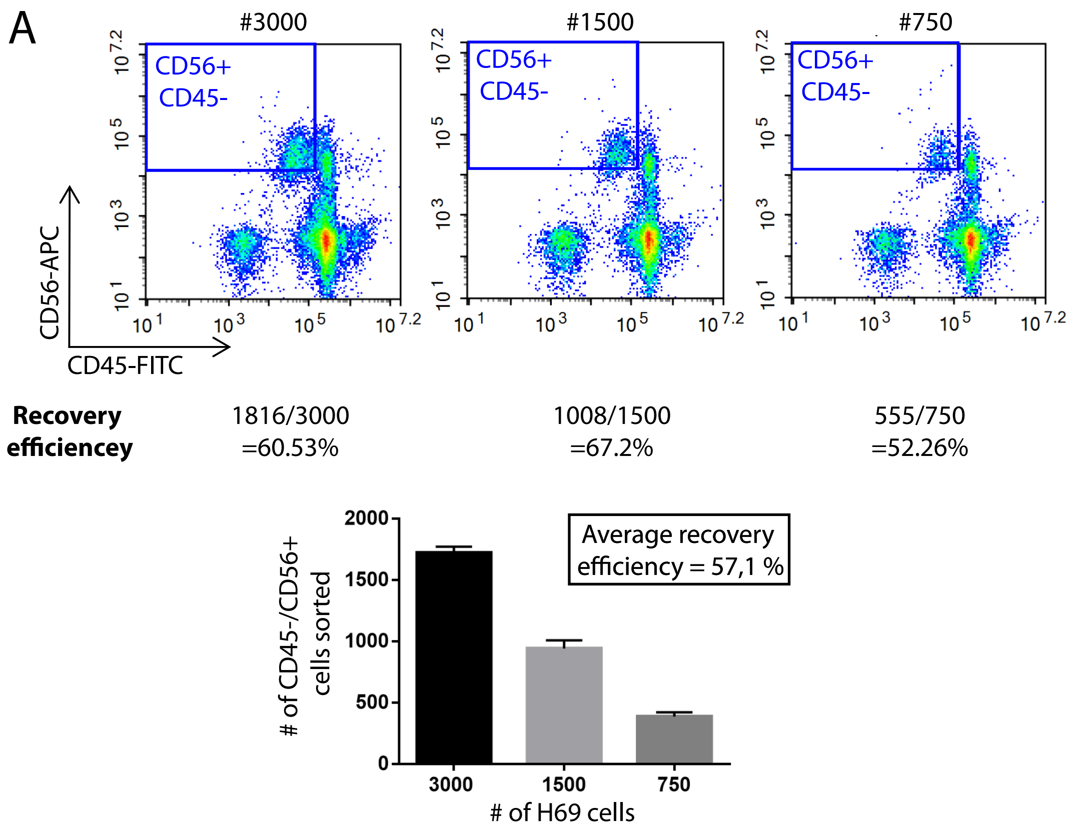


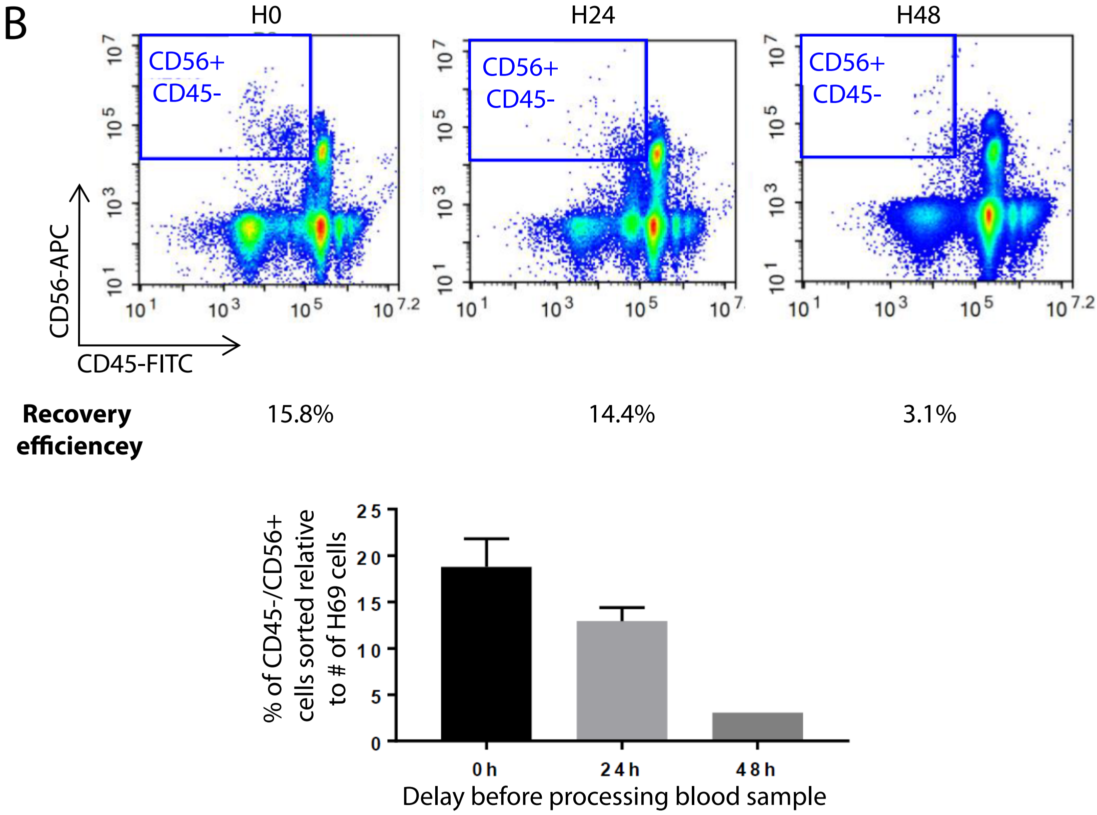

### Supplemental Fig.4

Supplemental Fig 4

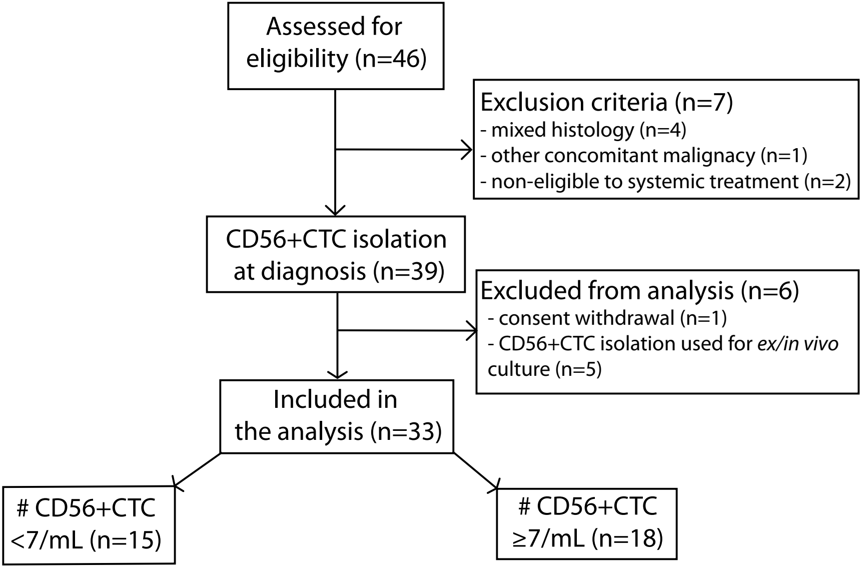

### Supplemental Fig.5

Supplemental Fig 5

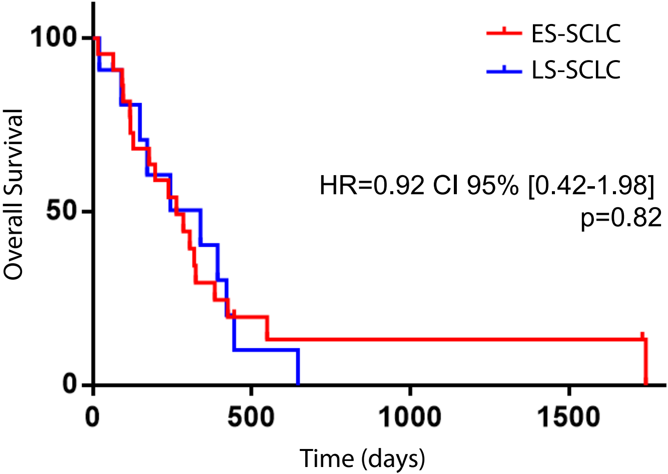
