## Supplemental Fig.3 for "Genomic characteristics and clinical significance of CD56+ Circulating Tumor Cells in Small Cell Lung Cancer"

### Supplemental Fig3

A

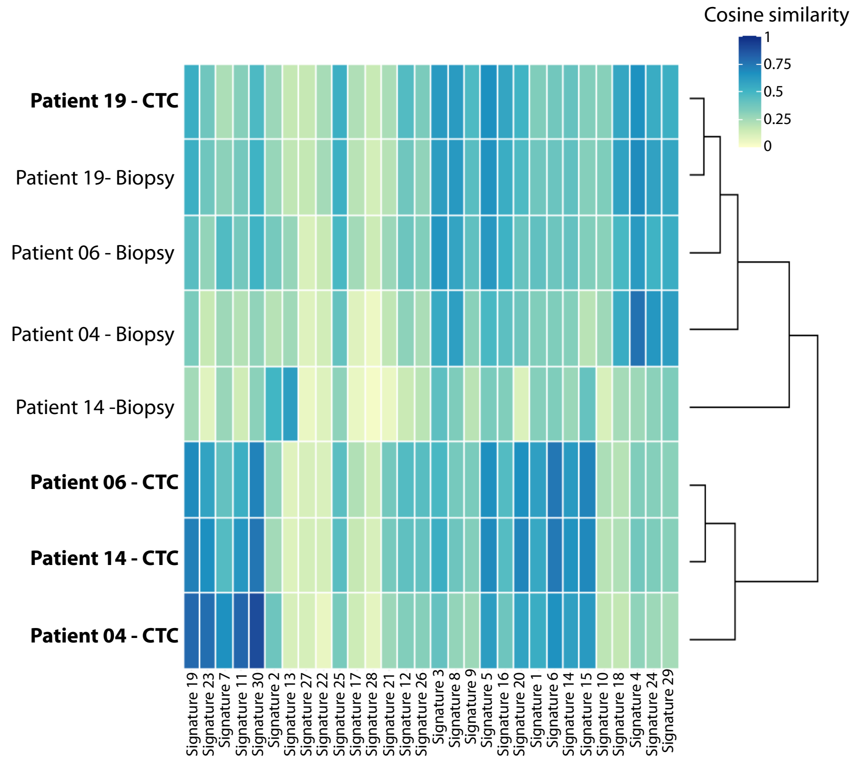


B

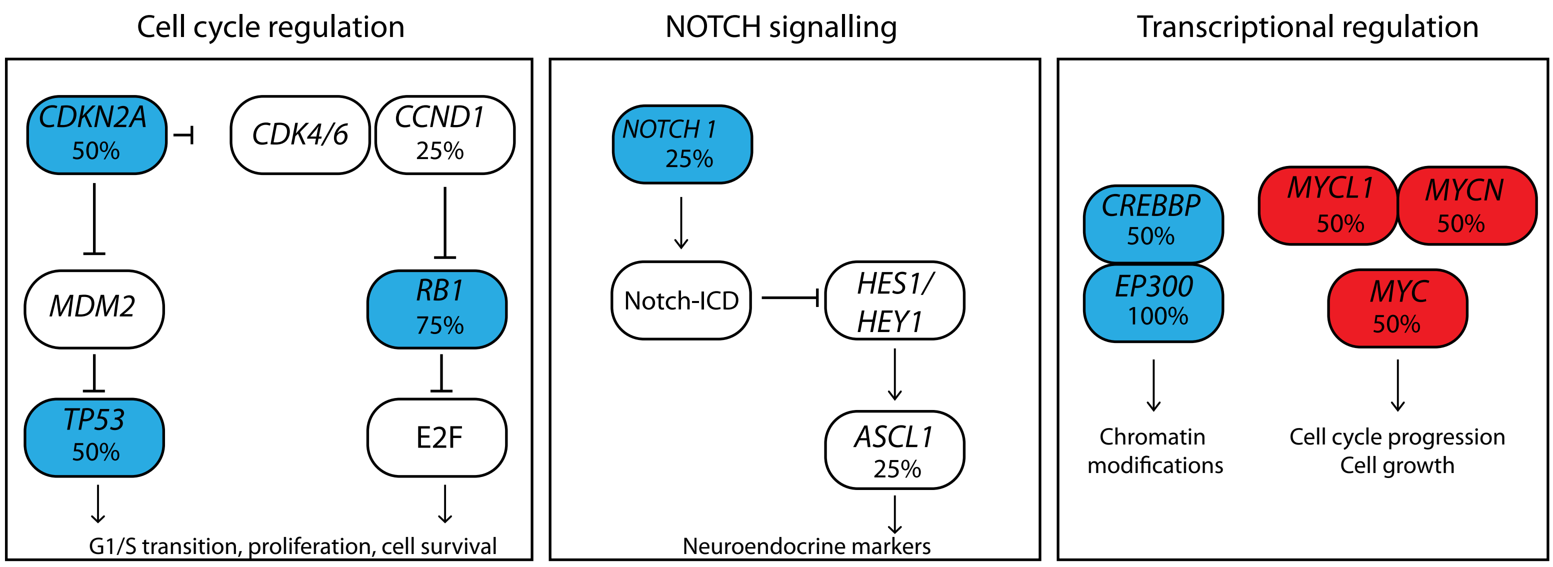


C

**CD56+CTC 75 hub genes**

| | | | | |
|---|---|---|---|---|
| AR | COL5A1 | HCK | LRP6 | PROX1 |
| ARHGAP35 | COL9A1 | HDAC6 | MAP3K12 | PTK7 |
| ARNT | COL9A3 | HIF1A | MDM2 | PTPRS |
| AXL | CREBBP | HIF3A | MECP2 | ROCK2 |
| BAX | CSF1 | HIPK2 | MED1 | RUNX2 |
| BMI1 | DNM2 | INSR | MEIS1 | SIX4 |
| BRAF | DSCAM | INSRR | MTOR | SMAD4 |
| BTRC | EGLN1 | ITGAV | NF1 | STAT3 |
| CACNA1A | EGR1 | ITGB1 | NR4A3 | STAT5B |
| CACNA1C | EIF2AK3 | JARID2 | NRG1 | TP53 |
| CHD3 | EP300 | KALRN | PARP1 | TRIM28 |
| CITED2 | EPHB1 | KDM5B | PIP5K1C | TSC1 |
| COL27A1 | ERBB2 | KMT2A | PLCB1 | WNT1 |
| COL4A2 | ESR1 | KMT2E | PRKCG | YAP1 |
| COL4A3 | FZD5 | LAMC3 | PRKDC | ZBTB7A |

**CD56+CTC 23 core hub genes**

| | | |
|---|---|---|
| AR | ERBB2 | NRG1 |
| ARHGAP35 | ESR1 | PARP1 |
| ARNT | HIF1A | SMAD4 |
| BAX | KMT2A | STAT3 |
| BRAF | MDM2 | STAT5B |
| CREBBP | MTOR | TP53 |
| DNM2 | NF1 | TSC1 |
| EP300 | NR4A3 | |

GENE NAME : common with all tissue databases

GENE NAME : common with 1 or 2 tissue databases

D

| Rank | Term | Definition | Enrichment | Pvalue |
|---|---|---|---|---|
| 1 | 1234158 | Regulation of gene expression by Hypoxia-inducible Factor | 4/11 | 0.0041 |
| 2 | 5663202 | Diseases of signal transduction | 12/386 | 0.0086 |
| 3 | 8864260 | Transcriptional regulation by the AP-2 (TFAP2) family of transcription factors | 4/38 | 0.014 |
| 4 | 6804114 | TP53 Regulates Transcription of Genes Involved in G2 Cell Cycle Arrest | 3/18 | 0.0162 |
| 5 | 6804760 | Regulation of TP53 Activity through Methylation | 3/19 | 0.0238 |
| 6 | 3108232 | SUMO E3 ligases SUMOylate target proteins | 7/181 | 0.0254 |
| 7 | 6803204 | TP53 Regulates Transcription of Genes Involved in Cytochrome C Release | 3/20 | 0.0295 |
| 8 | 2990846 | SUMOylation | 7/187 | 0.0352 |
| 9 | 8878159 | Transcriptional regulation by RUNX3 | 5/96 | 0.0374 |
| 10 | 2219528 | PI3K/AKT Signaling in Cancer | 5/102 | 0.0416 |
| 11 | 8941855 | RUNX3 regulates CDKN1A transcription | 2/7 | 0.0464 |
| 12 | 1234174 | Cellular response to hypoxia | 4/75 | 0.0499 |
