## Supplemental Table for "Genomic characteristics and clinical significance of CD56+ Circulating Tumor Cells in Small Cell Lung Cancer"

Table supplemental 1 :

| SCLC CD56+CTC gene signature | | | CGC.CommonGenes | IntOGen.Mutational.DriverGenes | MyCancerGenome.SCLC |
| --- | --- | --- | --- | --- | --- |
| AR | EP300 | MEIS1 | AR | AR | BRAF |
| ARHGAP35 | EPHB1 | MTOR | ARNT | ARHGAP35 | CREBBP |
| ARNT | ERBB2 | NF1 | BAX | BRAF | EP300 |
| AXL | ESR1 | NR4A3 | BRAF | CREBBP | ERBB2 |
| BAX | FZD5 | NRG1 | CREBBP | EP300 | MDM2 |
| BMI1 | HCK | PARP1 | DNM2 | ERBB2 | MTOR |
| BRAF | HDAC6 | PIP5K1C | EP300 | ESR1 | PARP1 |
| BTRC | HIF1A | PLCB1 | ERBB2 | KMT2A | TP53 |
| CACNA1A | HIF3A | PRKCG | ESR1 | MDM2 | TSC1 |
| CACNA1C | HIPK2 | PRKDC | HIF1A | MTOR |  |
| CHD3 | INSR | PROX1 | ITGAV | NF1 |  |
| CITED2 | INSRR | PTK7 | KMT2A | NRG1 |  |
| COL27A1 | ITGAV | PTPRS | MDM2 | SMAD4 |  |
| COL4A2 | ITGB1 | ROCK2 | MTOR | STAT3 |  |
| COL4A3 | JARID2 | RUNX2 | NF1 | STAT5B |  |
| COL5A1 | KALRN | SIX4 | NR4A3 | TP53 |  |
| COL9A1 | KDM5B | SMAD4 | NRG1 | TSC1 |  |
| COL9A3 | KMT2A | STAT3 | SMAD4 |  |  |
| CREBBP | KMT2E | STAT5B | STAT3 |  |  |
| CSF1 | LAMC3 | TP53 | STAT5B |  |  |
| DNM2 | LRP6 | TRIM28 | TP53 |  |  |
| DSCAM | MAP3K12 | TSC1 | TSC1 |  |  |
| EGLN1 | MDM2 | WNT1 |  |  |  |
| EGR1 | MECP2 | YAP1 |  |  |  |
| EIF2AK3 | MED1 | ZBTB7A |  |  |  |
